## Supplementary figures and images for "Temporally resolved early BMP-driven transcriptional cascade during human amnion specification"

### Figur e S2

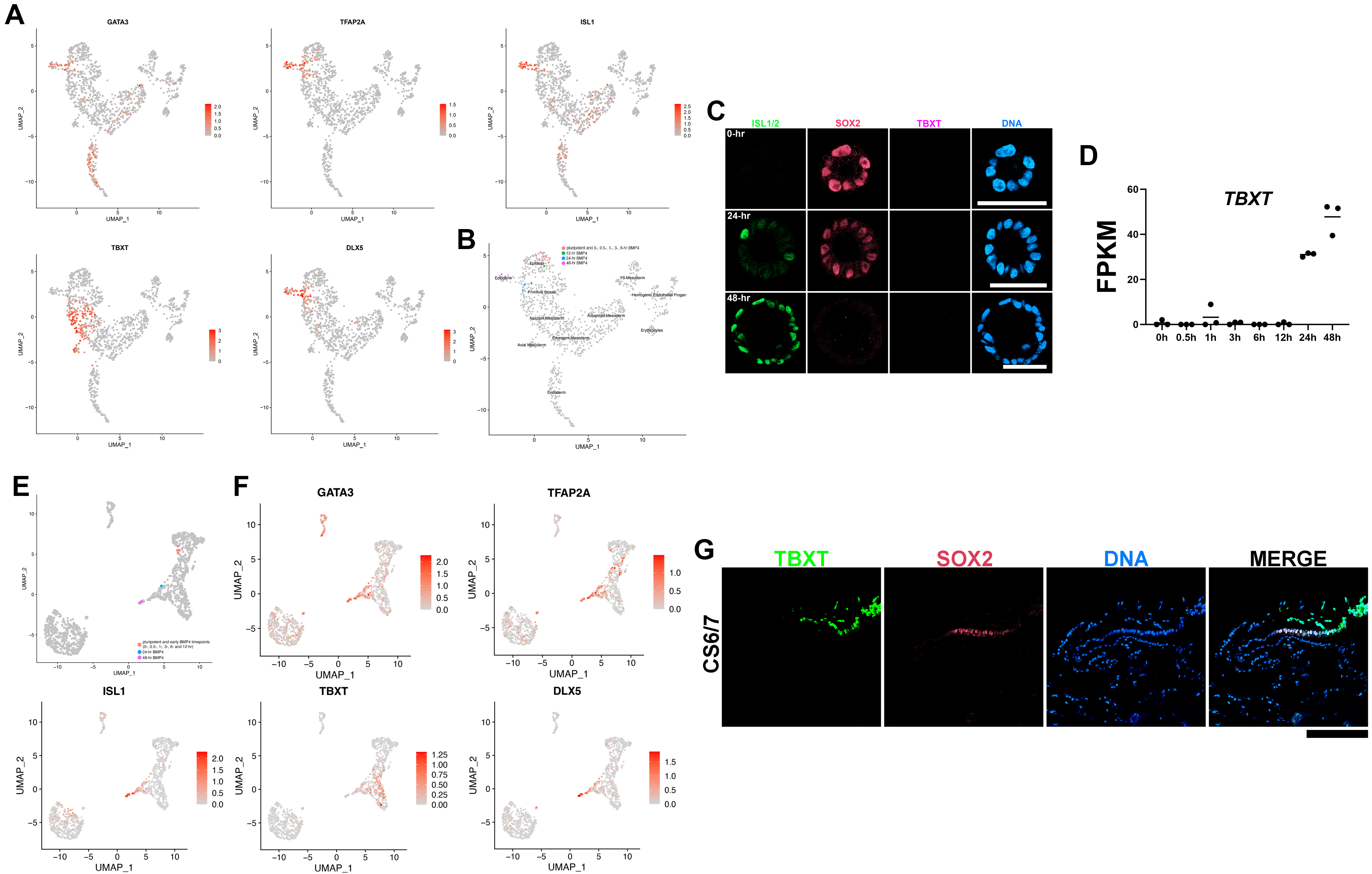

### Figure S1

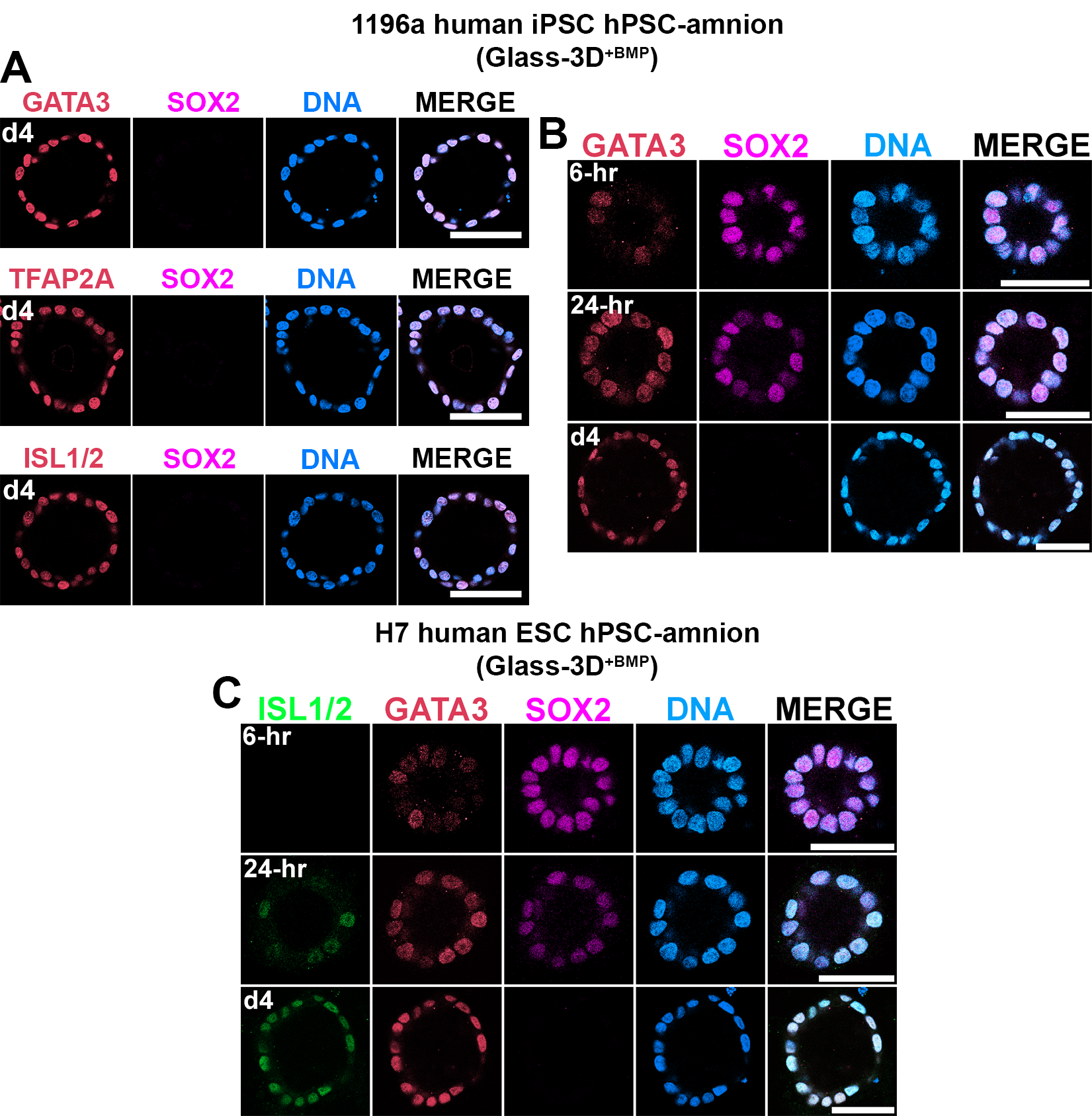

### Figure S3

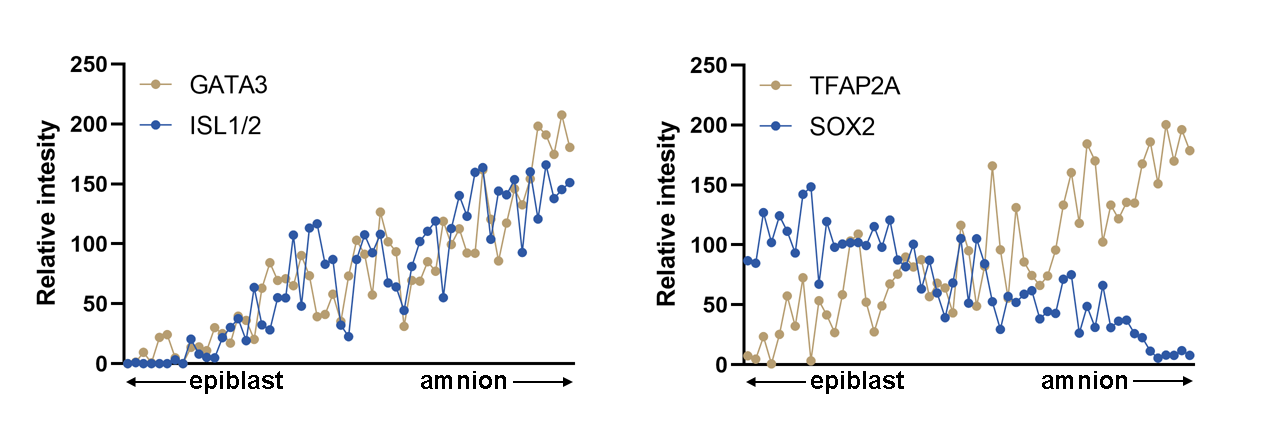

### Figure S4

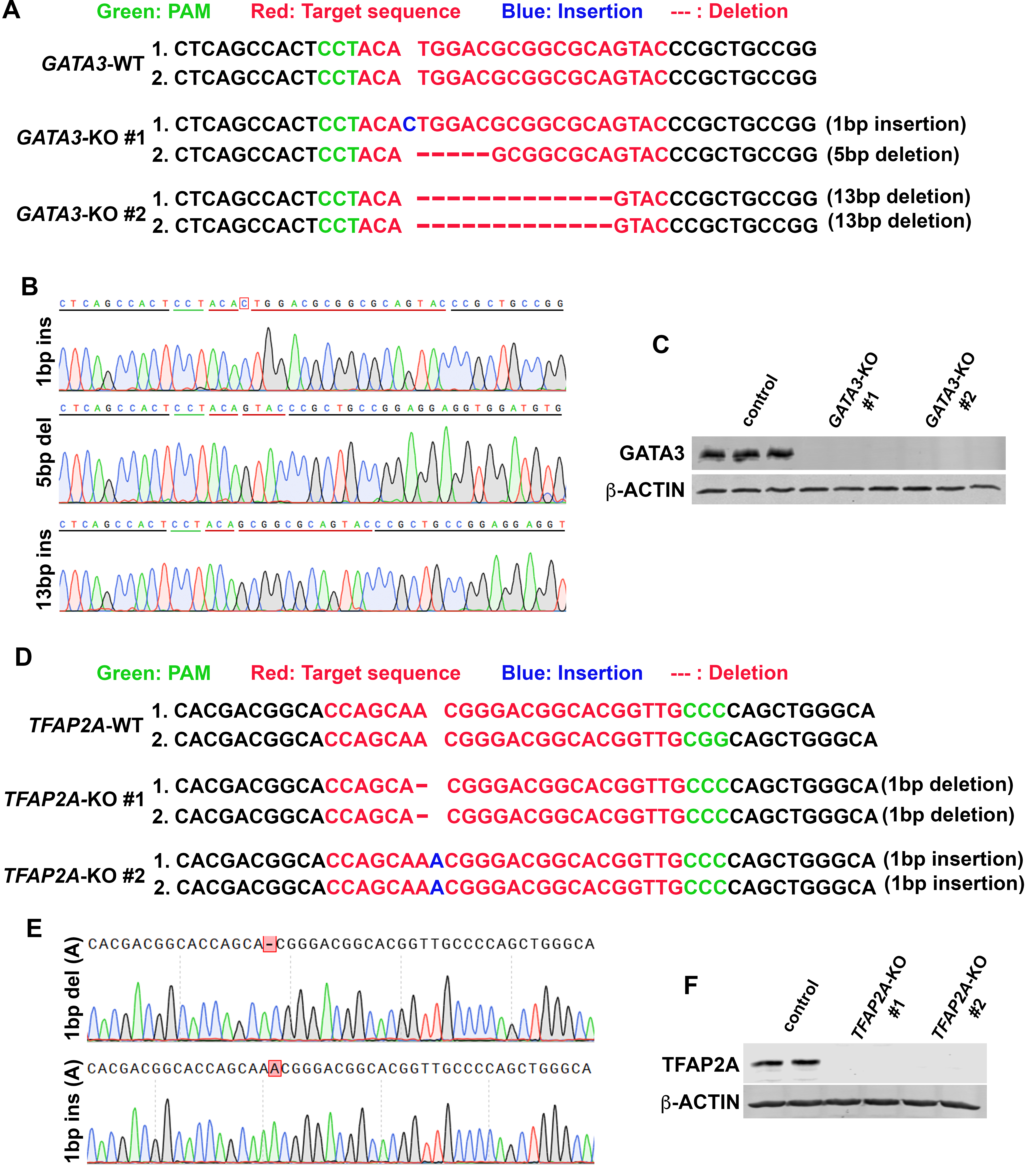

### Figure S5

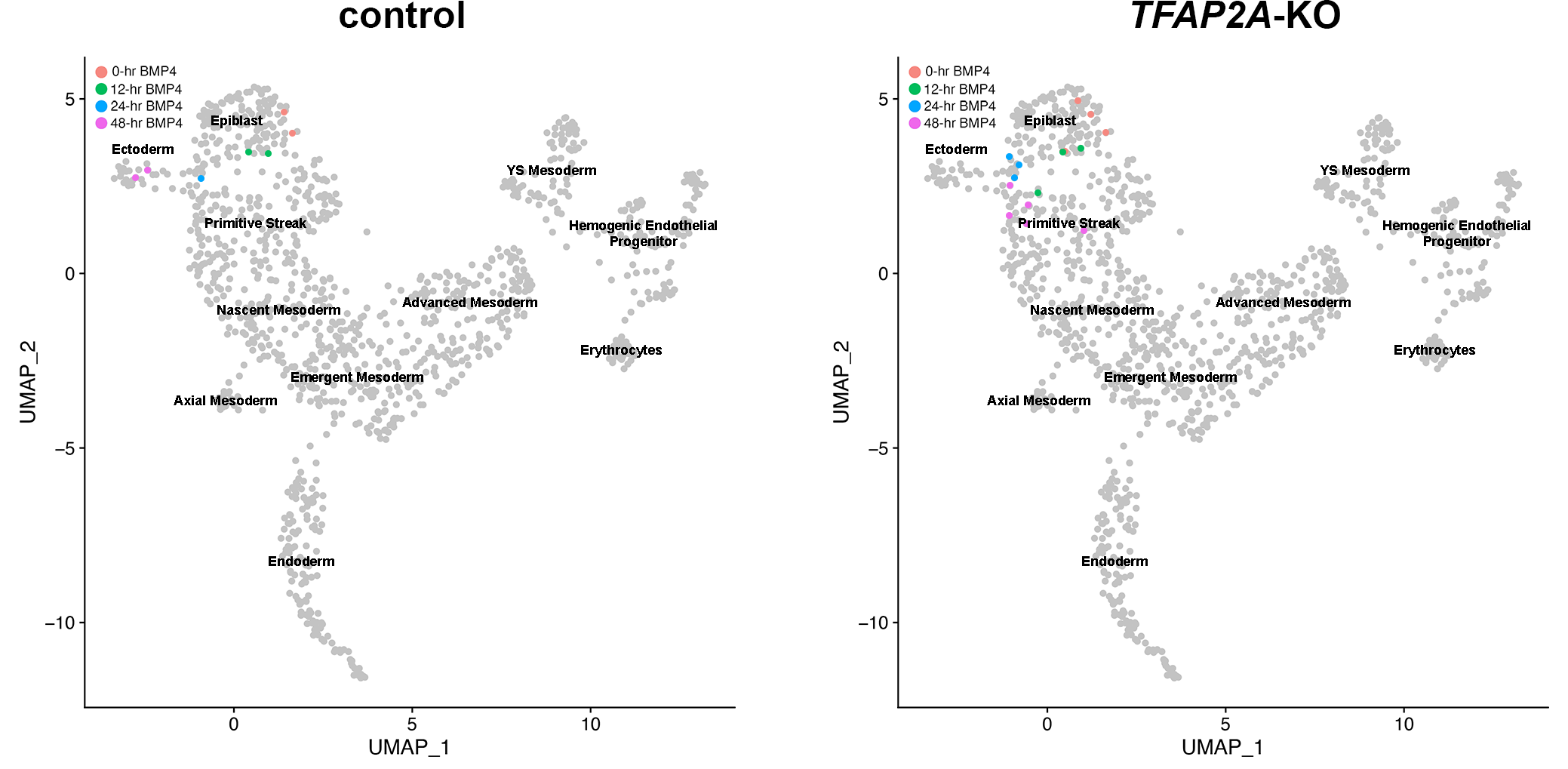
